# Proto-MHC signatures in the lamprey genome

**DOI:** 10.1101/2025.03.16.643586

**Authors:** Yoichi Sutoh, Kanako Ono, Shiori Minabe, Samadhi Senanayake, Hideki Ohmomo, Tsuyoshi Hachiya, Atsushi Shimizu, Masayuki Hirano

## Abstract

Jawless vertebrates, which diverged from jawed vertebrates approximately 550 million years ago, possess an adaptive immune system characterized by two distinct lineages of lymphocytes bearing unique antigen receptors. However, despite decades of research, the mechanism of antigen presentation in these organisms remains unclear. In this study, we report that chromosomes 22, 50, and 59 in the sea lamprey (*Petromyzon marinus*) exhibit synteny with the human major histocompatibility complex (MHC) region. The syntenic region spans from *DHX16* to *RING1* (chromosome 6: 30,517,301–33,034,069) in the human genome (T2T-CHM13v2.0). Similar to the human MHC region, *P. marinus* chromosome 59 harbors homologs of *DDX39B*, *NOTCH*, *PBX*, *TNF,* and *BRD* genes, yet notably lacks the core MHC class I, II, and III (complement) genes. Genotyping of three independent lampreys showed no signatures of balancing selection, a characteristic feature of the human MHC. Additionally, these lamprey chromosomes exhibit synteny with chromosome 9 in amphioxus (*Branchiostoma lanceolatum*), which contains conserved homologs that form its “proto-MHC.” These findings suggest that chromosomes 22, 50, and 59 in *P. marinus* are a proto-MHC or its paralog.

## Introduction

Jawless vertebrates (lampreys and hagfishes) diverged from the common ancestor of vertebrates approximately 550 million years ago (Mya) (Janvier 2015; Boehm 2025). Fossil records indicate that jawless vertebrates closely related to modern lampreys existed around 360 Mya during the Late Devonian period, suggesting that they have retained the ancestral body plan of early vertebrates (Gess, Coates, and Rubidge 2006). Immunological studies have demonstrated that jawless vertebrates exhibit adaptive immune responses, including allograft rejection (Perey et al. 1968; Hildemann and Thoenes 1969) and humoral responses (Finstad and Good 1964; Pollara et al. 1970; Linthicum and Hildemann 1970; Acton et al. 1969), as well as secondary immune responses similar to those of jawed vertebrates. Since the discovery of variable lymphocyte receptors (VLRs) (Pancer et al. 2004; 2005; Rogozin et al. 2007; Kasamatsu et al. 2010; J. Li et al. 2013; Das et al. 2023), research on the immune system of jawless vertebrates has advanced significantly.

However, the molecular mechanism underlying allograft rejection in jawless vertebrates remains unclear. In jawed vertebrates, allograft rejection is primarily mediated by mismatches in major histocompatibility complex (MHC) genes— particularly the “core” MHC genes, such as *HLA-A, -B, -C, -DR, -DQ,* and *-DP* in humans—between the host and graft. The primary function of core MHC genes is antigen presentation, and maintaining a diverse repertoire of alleles provides an evolutionary advantage by preventing pathogens from evading immune detection. This selective pressure is an example of balancing selection, a mechanism that preserves multiple alleles within a population (Takahata, Satta, and Klein 1992; Yasukochi and Satta 2013). Consequently, allelic diversity is a defining feature of core MHC genes, reflecting their critical role in immune defense. While several polymorphic genes have been identified in jawless vertebrates (Takaba et al. 2013; Haruta, Suzuki, and Kasahara 2006), no conclusive evidence has been found for the existence of core MHC genes in these species.

Another key feature of the MHC is its organization as a gene cluster enriched with immune-related genes. The human MHC consists of three classes—MHC class I, II, and III—whose gene composition has been evolutionarily conserved. A syntenic gene cluster homologous to the MHC has been identified in amphioxus (*Branchiostoma lanceolatum*), referred to as the “proto-MHC” (Abi-Rached et al. 2002; Veríssimo et al. 2023; Shiina et al. 2003). The proto-MHC is composed of evolutionarily conserved homologs of MHC-associated genes, including *RXRA/B/G, BAT1/DDX39, BRD2/3/4/T, C3/C4/C5, CACNA1A/B/E, NOTCH1/2/3/4, PSMB5/8, PSMB7/10,* and *PBX1/2/3/4,* but lacks the highly diverse core MHC genes.

In this study, we investigated the genomic regions homologous to the MHC in lampreys using the latest chromosome-level genome assembly and long-read sequencing technologies. Syntenic analysis revealed significant collinearity between several lamprey chromosomes and the human MHC region. Additionally, long-read whole-genome sequencing of three individual lampreys enabled the determination of allelic diversity at these loci. Our findings provide insights into the evolutionary history of the MHC in early vertebrates.

## Materials and Methods

### Animals

Sea lampreys (*Petromyzon marinus*) were captured in the Great Lakes and Maine, United States. Blood was collected from tail-severed lampreys and diluted with 0.67× PBS containing 30 mM EDTA, after the animals were sedated with Syncaine (MS-222, 100 mg/L, Syndel USA). Buffy coat leukocytes were collected by centrifugation at 400 × g for 15 minutes at room temperature. Genomic DNA was purified from the leukocytes using the DNeasy Blood & Tissue Kit (QIAGEN). All lamprey experiments were approved by the Institutional Animal Care and Use Committee at Emory University. Out of five DNA samples from individual sea lampreys, three with higher quality and sufficient quantity (*Pm1, Pm2,* and *Pm5*) were selected for sequencing.

### Sequencing, variant calling, and genotyping

A total of 1 µg of purified DNA was used for library preparation with the Ligation Sequencing DNA V14 kit (SQK-LSK114, Oxford Nanopore Technologies) and the NEBNext^®^ Companion Module for Oxford Nanopore Technologies^®^ Ligation Sequencing (NEB, E7180S), following the manufacturer’s protocols. The prepared library was applied to a PromethION Flow Cell (R10.4.1, FLO-PRO114M, Oxford Nanopore Technologies) and sequenced using the PromethION 2 Solo (Oxford Nanopore Technologies) for approximately 24 hours. Sequencing was monitored using MinKNOW (version 23.11.7).

Variant calling was performed using Dorado (version 0.9.1) in super accuracy mode with the basecalling model *dna_r10.4.1_e8.2_400bps_sup@v5.0.0*. The file format was converted from FASTQ to BAM using Samtools (version 1.21) (Danecek et al. 2021), after which the sequencing reads were mapped to the *P. marinus* reference genome (*kPetMar1.pri*, NCBI RefSeq assembly: GCF_010993605.1) using Minimap2 (version 2.26-r1175)(H. Li 2021). The mapped data was then sorted and used for variant calling with BCFtools (version 1.21)(Danecek et al. 2021).

### Nucleotide variation and selection analysis

Quality control was performed using BCFtools, excluding variants that met the following criteria: low quality (QUAL < 20), excessive depth (DP > twofold of mean depth), or missing count ≥2. Insertions and deletions (indels) were normalized and separated from single nucleotide polymorphisms (SNPs). Adjacent indels and SNPs within 5 bp or 2 bp, respectively, were excluded.

For comparison, publicly available human genome variation data were retrieved from the Human Genome Structural Variation Consortium, Phase 3 (Logsdon et al. 2024), which includes 65 individuals of diverse ancestries. Variants with a call rate below 0.67 were excluded, and adjacent indels and SNPs within 5 bp or 2 bp, respectively, were removed.

Nucleotide diversity and Tajima’s D (Tajima 1989) were calculated for every 10 kb using VCFtools (version 0.1.16)(Danecek et al. 2011) based on SNP data. The results were plotted using R (version 4.3.2, released 2023-10-31) and RStudio (version 2023.12.1+402) with ggExtra package (version 0.10.1).

### Synteny analysis

Synteny analysis was performed using the MCScanX toolkit (Wang et al. 2024) for multiple collinearity scanning. MCScanX requires General Feature Format (GFF) files and protein sequence (FASTA) files for genome assembly. These files were retrieved from the latest sea lamprey (*P. marinus*), human (*Homo sapience*) and amphioxus (*B. lanceolatum*) genome assemblies: *kPetMar1.pri* (NCBI RefSeq assembly: GCF_010993605.1), *T2T-CHM13v2.0* (NCBI RefSeq assembly: GCF_009914755.1) (Nurk et al. 2022), and *klBraLanc5.hap2* (NCBI RefSeq assembly: GCF_035083965.1), respectively.

Homology searches between human and lamprey protein sequences were conducted using DIAMOND (Buchfink, Reuter, and Drost 2021). Human and lamprey protein FASTA files were combined and used as both the database and query. The homology search was performed using BLASTP mode with the ultra-sensitive option, and alignments with *e*-values ≤ 1e-10 were retained. The lamprey and human GFF files were also combined and reformatted according to MCScanX documentation.

The MCScanX program was executed using the input data (BLAST results and reformatted GFF file) with loose gap penalty settings (*max gaps* = 200, *gap penalty* = 0), considering the evolutionary divergence between vertebrate classes. The output (collinearity file) were visualized using SynVisio (Bandi and Gutwin 2020) and AccuSyn (Nunez Siri et al. 2020).

Orthogroups for proto-MHC were identified using OrthoFinder (version 2.5.5) (Emms and Kelly 2019) with default settings on Docker Desktop (version 4.37.2 [179585]). Protein sequences from the latest genome assemblies of human (*T2T-CHM13v2.0*), zebrafish (*GRCz11*, NCBI RefSeq assembly: GCF_000002035.6), shark (*sHemOce1.pat.X.cur,* NCBI RefSeq assembly: GCF_020745735.1), lamprey (*kPetMar1.pri*), and amphioxus (*klBraLanc5.hap2*) were used for the analysis.

### Data availability

The long-read sequencing data from individual sea lamprey genomes are available under NCBI BioProject accession **PRJNA1228496**. Sample codes for the methods used in this study are available on GitHub: https://github.com/yyoi/Lamprey-20250210

## Results

### Lamprey chromosomes showing synteny with the human MHC

The human extended MHC (xMHC) region is defined as spanning from *SCGN* to *SYNGAP1* (Horton et al. 2004), corresponding to positions 25,518,025–33,275,052 on chromosome 6 in the latest human genome assembly (*T2T-CHM13v2.0*) (Nurk et al. 2022).

Synteny analysis of the latest lamprey genome assembly (*kPetMar1.pri*) was conducted using the MCScanX toolkit, which identifies collinear blocks of homologous genes based on protein sequence similarity (Wang et al. 2024). This analysis revealed that 14 lamprey chromosomes contain regions with significant synteny (e-value < 1e-5) to human chromosome 6 (Figure 1A). Among these, synteny in chromosomes 50 and 59 specifically corresponded to the xMHC region.

**Figure 1.**
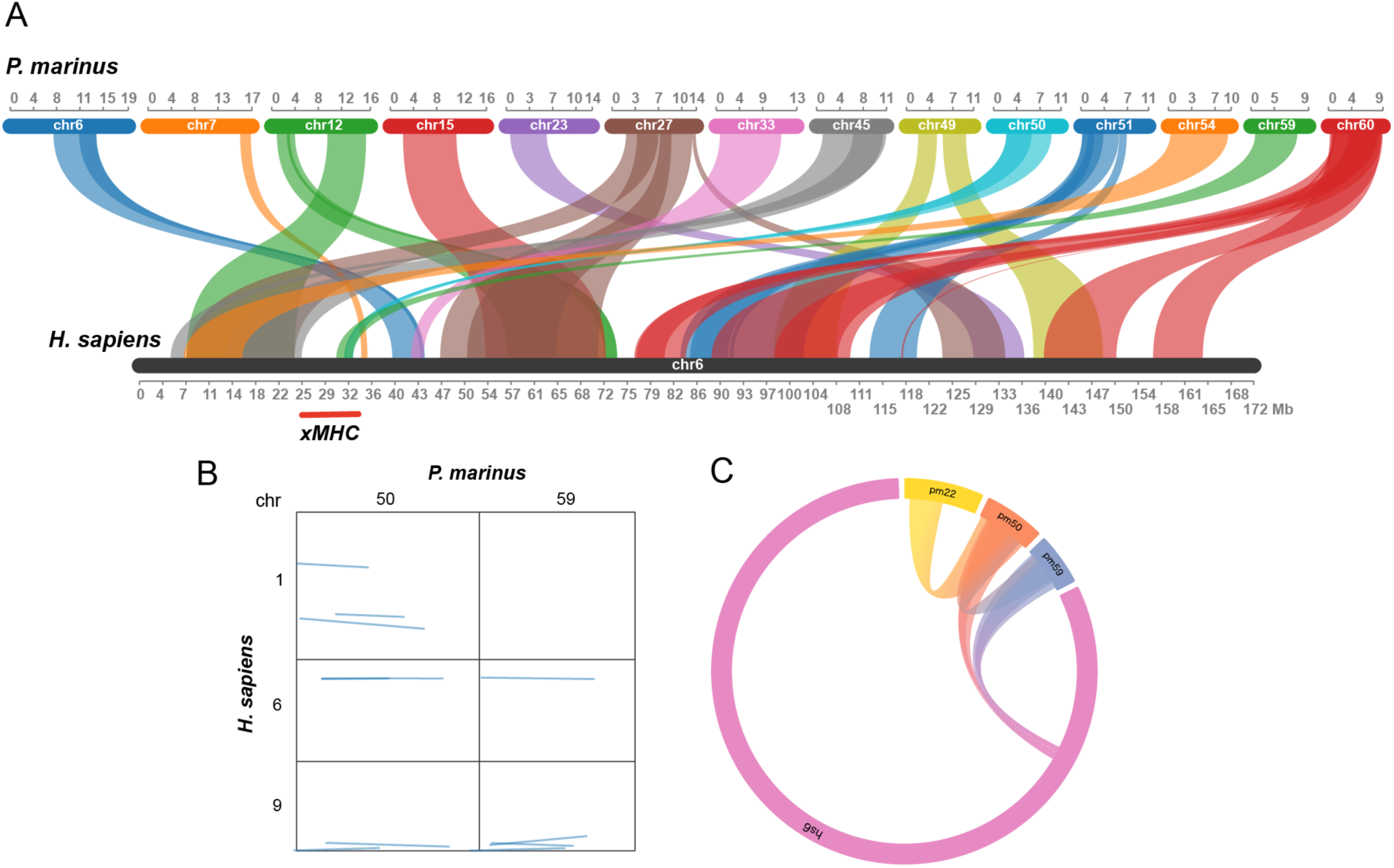
Synteny between human and lamprey chromosomes. Collinearity blocks in the sea lamprey (*Petromyzon marinus*) and human (*Homo sapiens*) genomes were identified using the MCScanX toolkit and visualized with SynVisio (A, B). Synteny with human chromosome 6 (shown in black) was identified across 14 lamprey chromosomes, with chromosomes 50 (light blue) and 59 (green) specifically encompassing the human extended MHC (xMHC) region. (A). A dot plot comparison between human and lamprey chromosomes (B) shows that synteny spans only a narrow region of human chromosome 6 but extends across nearly the entire length of the corresponding lamprey chromosomes. Inter- and intra-genome synteny analysis, visualized with AccuSyn, highlights collinearity blocks among lamprey chromosomes 22 (yellow), 50 (red), and 59 (blue), but not human chromosome 6 (purple) (C). The whole MCScanX collinearity output file is provided as *Supplementary Data 1*. *Pm*, *Petromyzon marinus*; *Hs*, *Homo sapiens*; *chr*, chromosome.

The relative sizes of the syntenic regions was “asymmetrical” between humans and lampreys: in humans, synteny covered only a small portion of chromosome 6, whereas in lampreys, nearly the entire length of chromosomes 50 and 59 exhibited synteny with the human xMHC (Figure 1B). Additionally, chromosome 50 showed synteny with human chromosomes 1 and 9, while chromosome 59 showed synteny with chromosome 9. Furthermore, lamprey chromosomes 50 and 59 exhibited intra-synteny with lamprey chromosome 22 (Figure 1C), although the synteny between lamprey chromosome 22 and human chromosome 6 was not statistically significant.

The syntenic regions in chromosomes 50 and 59, defined as spanning from the first to the last gene in collinearity, overlapped with the xMHC region (Table 1).

**Table 1.** Significant collinearity blocks with human chromosome 6.

| Lamprey |  | Human |  | MCScanX |  |
| --- | --- | --- | --- | --- | --- |
| chr | region<sup>a</sup> | chr | region<sup>a</sup> | e-value | direction |
| 50 | 2,084,530..9,367,401 | 6 | 31,716,432..32,016,865 | 0 | + |
| 50 | 2,069,471..6,307,142 | 6 | 33,098,485..31,962,530 | 0 | - |
| 59 | 711,047..7,498,040 | 6 | 30,517,301..33,034,069 | 9.40E-08 | + |

a, region from the beginning of the first gene to the end of the last gene in the collinearly. chr, chromosome.

The syntenic regions differed between the two lamprey chromosomes (Figure 2; Supplementary Data 2 for registered gene names in the NCBI Gene database). While chromosome 50 encompassed the region from the end of MHC class III to class II, chromosome 59 spanned the middle of MHC class I, class III, and class II. Both chromosomes contained homologs of genes characteristic of the MHC.

**Figure 2.**
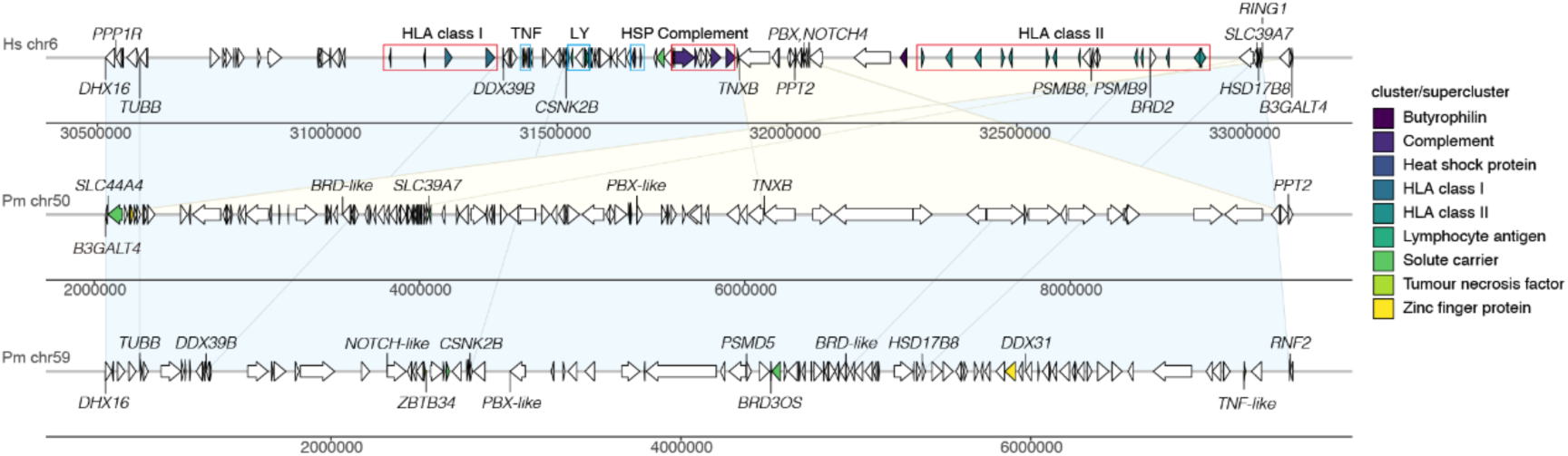
Genetic map of the syntenic regions. The syntenic regions identified in lamprey chromosomes 50 and 59 are represented on a genetic map for comparison. Cluster and supercluster classifications are based on Horton et al. (Horton et al. 2004). Background colors indicate syntenic regions: light yellow for chromosome 50 and light blue for chromosome 59. See *Supplementary Data 2* for the registered gene names in the NCBI Gene database. *Pm*, *Petromyzon marinus*; *Hs*, *Homo sapiens*; chr, chromosome.

Lamprey chromosome 50 harbored solute carrier proteins (*SLC44A4, SLC39A7*), bromodomain-containing protein 4-like (*BRD4-like* in Figure 2), pre-B-cell leukemia transcription factor 1-like (*PBX1-like*), and palmitoyl-protein thioesterase 2 (*PPT2*). Similarly, chromosome 59 contained DEAH-box helicases (*DHX16, DHX31*), spliceosome RNA helicase DDX39B (*DDX39B*), neurogenic locus notch homolog protein 1-like (“*NOTCH1-like*” in Figure 2), zinc finger proteins (*ZBTB34, LOC116955355*), and a tumor necrosis factor-like gene (*TNF-like*).

However, no homologs of core MHC genes, such as *HLA-A, -B, -C, -DR, -DQ*, or *-DP*, were identified in the lamprey chromosomes. Additionally, no complement genes were found in these regions.

### Diversity and signatures of selection in the lamprey genome

To identify specific genomic sites under balancing selection, a selective force that maintains allelic diversity, we obtained draft whole-genome sequencing data from three individual lampreys using Nanopore sequencing (Table 2).

**Table 2.**
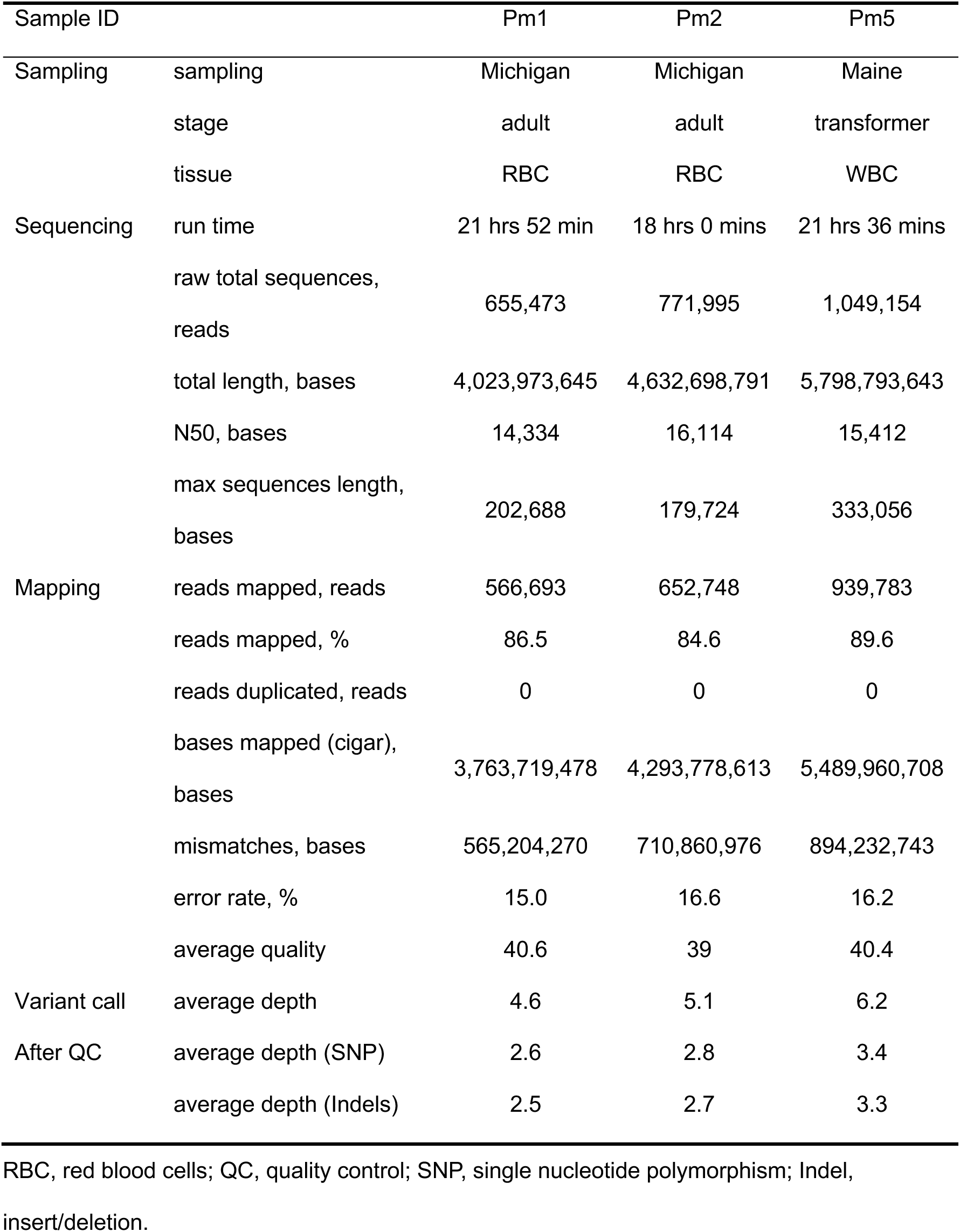
Summary of sequencing and genotyping for three individual lampreys.

By mapping the sequencing data to the reference genome, we identified 21,813,065 single nucleotide polymorphisms (SNPs) and 2,934,138 insertions/deletions (indels). After quality control filtering, 11,516,860 SNPs and 1,545,394 indels remained. Using this dataset, we calculated two genomic metrics: nucleotide diversity (π) as a measure of genetic variation and Tajima’s D as an indicator of selection pressure (Figure 3).

**Figure 3.**
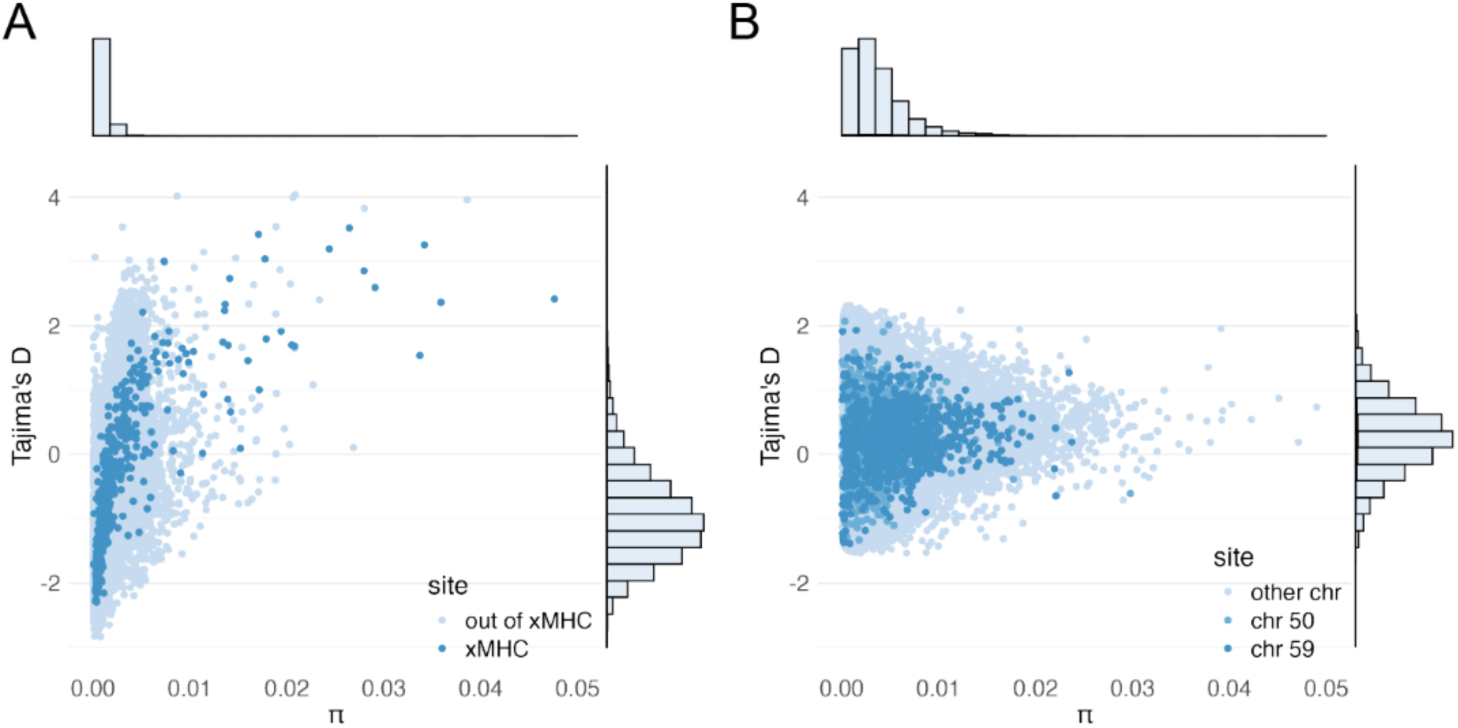
Genetic variation and selection in three lampreys. Nucleotide diversity (π) and Tajima’s D statistics, which indicate selection trends, were estimated using a 10 kb sliding window across the genome. The xMHC region (dark blue) exhibited high π and Tajima’s D values (A). However, no corresponding signature of balancing selection was observed at any site in the lamprey genome, including regions in chromosome 50 (blue) and chromosome 59 (dark blue) (B).

The xMHC region exhibited high nucleotide diversity and a positive Tajima’s D, a characteristic signature of balancing selection. In contrast, no genomic sites with a similar signature were identified in the three lamprey genomes, although some regions showed high nucleotide diversity.

### Synteny with “proto-MHC”

We compared lamprey chromosomes 22, 50, and 59 to the amphioxus (*Branchiostoma lanceolatum*) genome, which has been proposed to contain a proto-MHC in previous studies (Abi-Rached et al. 2002; Veríssimo et al. 2023; Shiina et al. 2003) (Figure 4). Lamprey chromosomes 22, 50, and 59 exhibited significant synteny with amphioxus chromosomes 9 and 14 (Figure 4A). However, the specific regions of collinearity varied among the lamprey chromosomes: chromosome 50 showed synteny with the forward region of amphioxus chromosome 9, while chromosomes 22 and 50 were syntenic with the backward region of chromosome 9 (Figure 4B). Additionally, only chromosome 22 displayed synteny with amphioxus chromosome 14.

**Figure 4.**
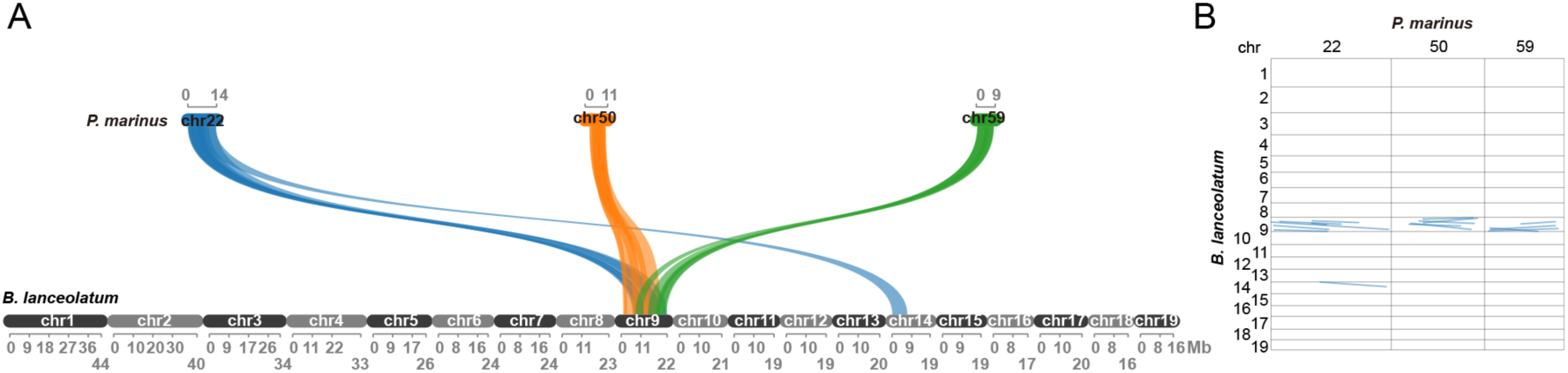
Synterny analysis with amphioxus genome. Collinearity blocks between the sea lamprey (*Petromyzon marinus*) and amphioxus (*Branchiostoma lanceolatum*) genomes were identified using the MCScanX toolkit and visualized with SynVisio (A, B). The full MCScanX collinearity output file is provided as Supplementary Data 3. See the Figure 1 legend for further details.

Lamprey chromosomes 22, 50, and 59 exhibited multiple overlapping collinear blocks with amphioxus chromosome 9, showing clearer synteny than with human chromosome 6 (Table 3).

**Table 3.** Significant collinearity blocks with lamprey and amphioxus chromosomes.

| Lamprey |  | Amphioxus |  | MCScanX |  |
| --- | --- | --- | --- | --- | --- |
| chr | region<sup>a</sup> | chr | region<sup>a</sup> | e-value | direction |
| 22 | 63,219..13,841,685 | 9 | 7,647,897..19,154,012 | 4.80E-10 | + |
| 22 | 354,447..6,850,535 | 9 | 12,928,450..19,212,884 | 0 | + |
| 22 | 1,073,014..8,250,744 | 9 | 6,652,025..10,998,141 | 1.40E-08 | + |
| 22 | 439,812..6,671,963 | 9 | 19,869,004..22,245,816 | 6.20E-06 | + |
| 22 | 4,869,779..10,427,752 | 9 | 5,736,867..8,493,194 | 0 | + |
| 22 | 4,786,262..6,577,428 | 9 | 9,790,306..11,551,123 | 0 | + |
| 22 | 5,803,869..13,550,631 | 14 | 196,217..7,392,062 | 3.50E-07 | + |
| 50 | 2,304,772..9,367,401 | 9 | 9,471,276..19,212,884 | 1.30E-19 | + |
| 50 | 2,084,530..8,152,896 | 9 | 11,141,914..13,647,549 | 4.60E-08 | + |
| 50 | 3,241,806..9,752,488 | 9 | 5,949,505..9,810,647 | 0 | + |
| 50 | 3,716,671..9,592,638 | 9 | 2,989,802..1,156,962 | 0 | - |
| 50 | 3,744,407..10,135,435 | 9 | 8,748,999..1,709,134 | 0 | - |
| 59 | 702,842..6,306,084 | 9 | 17,036,307..22,180,980 | 5.70E-08 | + |
| 59 | 711,047..8,334,447 | 9 | 19,212,149..13,208,203 | 4.00E-07 | - |
| 59 | 521,566..8,652,544 | 9 | 21,505,468..17,868,308 | 7.60E-08 | - |
| 59 | 4,275,713..8,318,024 | 9 | 10,560,715..7,050,220 | 8.80E-09 | - |

a, region from the beginning of the first gene to the end of the last gene in the collinearly.
chr, chromosome.

Finally, we identified homologs characteristic of the proto-MHC in lamprey and amphioxus chromosomes (Table 4). Amphioxus chromosome 9 contained most of the genes previously reported as proto-MHC components. In lamprey, four homologs were found on both chromosomes 22 and 50, while five homologs were identified on chromosome 59. A *PBX* homolog was conserved across all three lamprey chromosomes; however, complement genes (C3, C4, C5) were not present in these chromosomes. No proto-MHC homologs were identified on amphioxus chromosome 14. OrthoFinder confirmed most homologs as “orthogroups,” which are groups of genes descended from a single gene in the last common ancestor of a given set of species (Emms and Kelly 2019), except for *BRD2/3/4/T* (Table 4; see *Supplementary Data 4* for full orthogroup results).

**Table 4.**
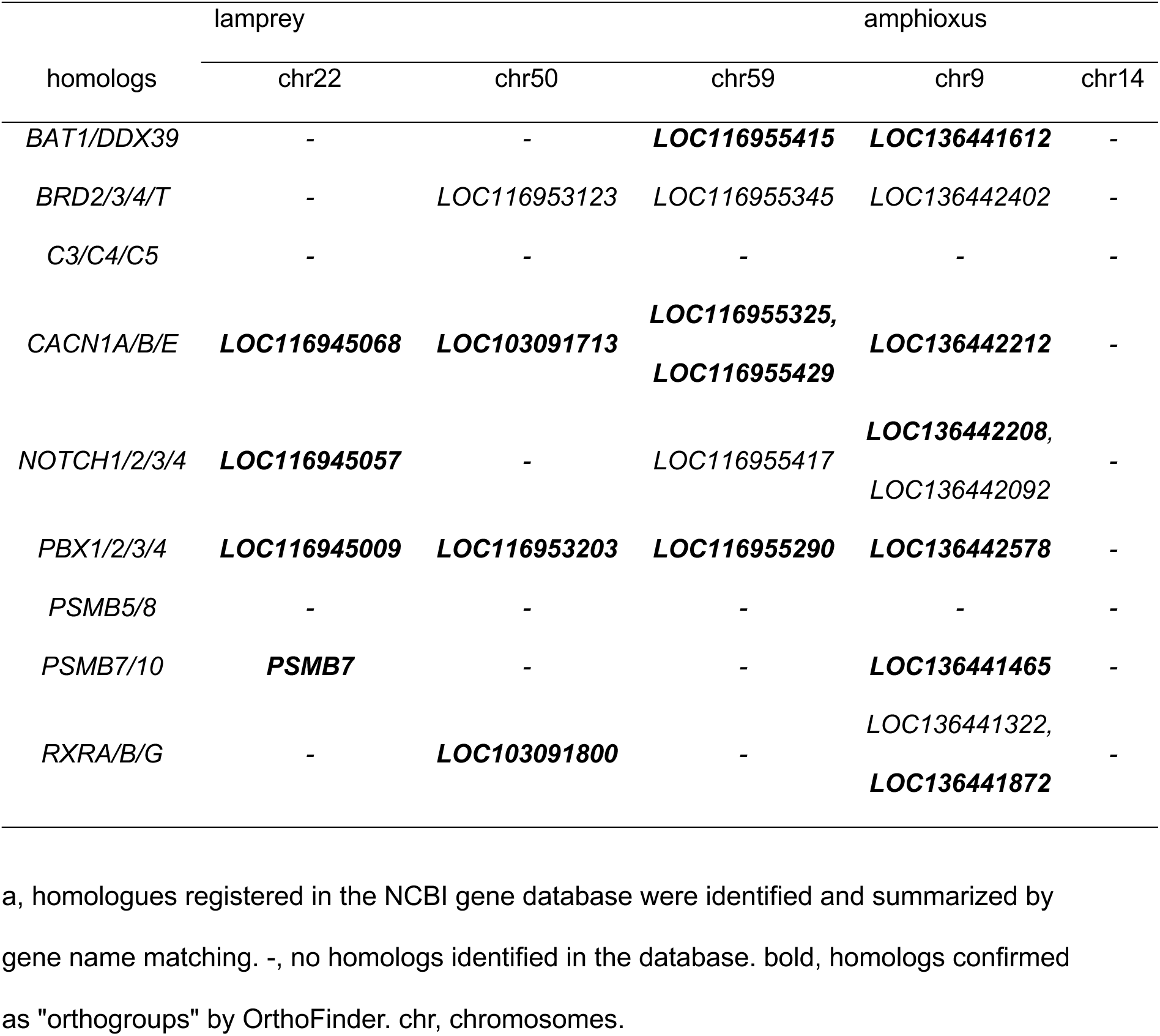
Conservation of gene sets characteristic of the proto-MHC^a^.

## Discussion

The existence of an MHC in jawless vertebrates has long been an unresolved question. A previous study based on the *Pmar_germline 1.0* assembly (GenBank: GCA_002833325.1) suggested that the sea lamprey possesses genomic regions homologous to the chicken MHC (Smith et al. 2018). However, a detailed analysis based on a chromosome-level assembly of the lamprey genome had not been conducted. The present study provides novel insights into the synteny of the MHC region in the lamprey genome.

The syntenic regions in lamprey chromosomes 50 and 59 encompassed MHC class I, II, and III regions but lacked core MHC genes. Furthermore, no clear signatures of balancing selection, as indicated by positive Tajima’s D values, were detected in the three lamprey genomes. These results do not support the presence of a conventional MHC with highly diverse alleles in jawless vertebrates.

Additionally, our analysis revealed significant synteny between lamprey chromosomes 50 and 59 and amphioxus chromosome 9 in the latest genome assembly (*klBraLanc5.hap2*). Amphioxus chromosome 9 retains nearly all homologs of the proto-MHC, as previously reported based on cosmid sequences (Shiina et al. 2003; Abi-Rached et al. 2002). The conservation of proto-MHC homologs suggests that chromosome 9 harbors the proto-MHC region or its paralogs. Moreover, the synteny between amphioxus chromosome 9 and lamprey chromosomes 22, 50, and 59, along with the distribution of their orthogroups, further supports the idea that these lamprey chromosomes contain proto-MHC regions rather than a conventional MHC, as seen in jawed vertebrates (Figure 5).

**Figure 5.**
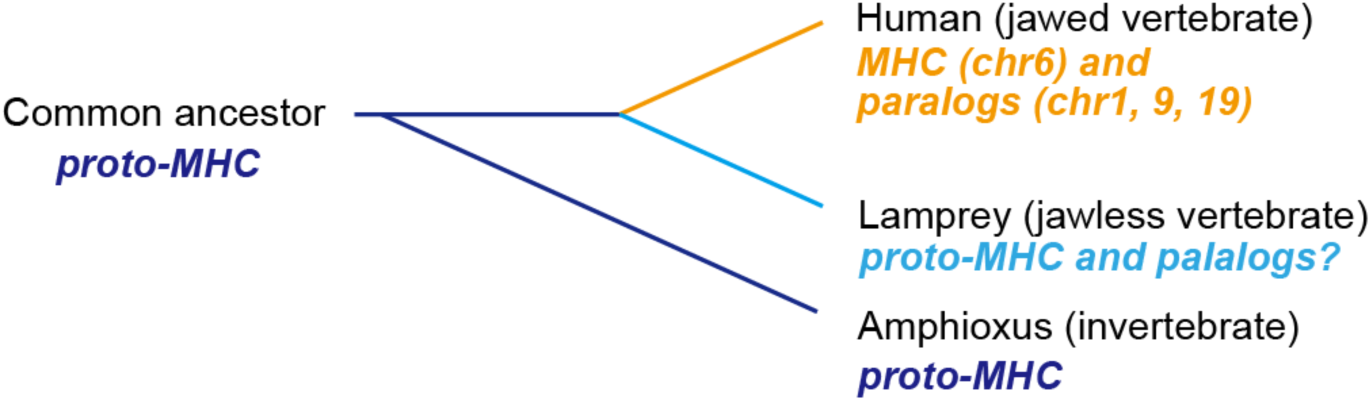
Schematic diagram of MHC evolution. The evolutionary stages of the MHC are represented within the animal phylogeny. Lamprey chromosomes retain proto-MHC signatures across multiple chromosomes but lack evidence of a conventional MHC with diverse alleles. The finding indicates that the emergence of a highly polymorphic MHC with diverse alleles likely occurred after the divergence of jawed vertebrates.

Genomic analyses of lampreys have suggested the presence of unique and complex duplication events in the jawless vertebrate lineage, including inter-chromosomal duplications (Timoshevskaya et al. 2023; Mehta et al. 2013; Zhu et al. 2021). In particular, lamprey chromosome 22 exhibited weaker similarity to the human MHC region, suggesting that it has undergone distinct duplication events compared to chromosomes 50 and 59 over evolutionary time.

This study has several limitations. First, the analysis relies on the current genome assembly and annotation, which may be subject to revision with future updates. Additionally, the lamprey polymorphism analysis was based on only three individuals from a limited geographical area, with relatively low sequencing coverage. To improve accuracy, future studies should include a larger sample size collected from a broader range of populations, with higher-coverage genomic data.

In conclusion, the sea lamprey genome exhibited proto-MHC signatures rather than characteristics of a conventional MHC in the present study. These findings suggest that the emergence of an MHC with allelic diversity, composed of MHC class I, II, and III genes, likely occurred after the divergence of jawed vertebrates from the common ancestor of vertebrates.

## Acknowledgement

We thank Miyuki Horie and Minako Chiba for their contributions to long-read sequencing. We also thank Shohei Komaki, Motoki Nakao, and Yayoi-Otsuka Yamasaki for their valuable advice on data analysis and manuscript writing.

## Author contribution

Y.S. designed the study and wrote the manuscript. S.S. and M.H. provided *P. marinus* genomic DNA. K.O. and S.M. conducted sequencing and primary data analysis. H.O., T.H., and A.S. provided critical insights as experts in bioinformatics and whole-genome analysis. M.H. provided critical insights as experts in evolutionary immunology.

